# Multiple origins of insular woodiness on the Canary Islands are consistent with palaeoclimatic aridification

**DOI:** 10.1101/2020.05.09.084582

**Authors:** Alexander Hooft van Huysduynen, Steven Janssens, Vincent Merckx, Rutger Vos, Luis Valente, Alexander Zizka, Maximilian Larter, Betül Karabayir, Daphne Maaskant, Youri Witmer, José Maria Fernández-Palacios, Lea de Nascimento, Ruth Molina Jaén, Juli Caujapé Castells, Águedo Marrero-Rodríguez, Marcelino del Arco, Frederic Lens

**Author notes:** corresponding author: Frederic Lens.

## Abstract

**Aim:** Insular woodiness, referring to the evolutionary transition from herbaceousness towards woodiness on islands, has arisen at least 38 times on the Canary Islands. Distribution patterns and physiological experiments have suggested a link between insular woodiness and increased drought stress resistance in current-day species, but we do not know in which palaeoclimatic conditions these insular woody lineages originated. Therefore, we estimated the timing of colonisation events and origin of woodiness of multiple Canary Island lineages and reviewed the palaeoclimate based on literature.

**Location:** Canary Islands (Spain).

**Taxon:** 37 lineages, including 24 insular woody and 13 non-insular woody (i.e. herbaceous, ancestrally woody, and derived woody).

**Methods:** To enable a simultaneous dating analysis for all 37 lineages, two chloroplast markers (*matK* and *rbcL*) for 135 Canary Island species and 103 closely related continental relatives were sequenced and aligned to an existing *matK-rbcL* dataset including ca 24,000 species that was calibrated with 42 fossils from outside the Canaries. After constraining the species to the family level, 200 RAxML runs were performed and dated with TreePL.

**Results:** Woodiness in 80-90% of the insular woody lineages originated within the last 7 Myr, coinciding with the onset of major aridification events nearby the Canaries (start of north African desertification, followed by Messinian salinity crisis); in ca 55-65% of the insular woody lineages studied, woodiness developed within the last 3.2 Myr during which Mediterranean seasonality (yearly summer droughts) became established on the Canaries, followed by dry Pleistocene glacial fluctuations.

**Main conclusions:** Although details of the initial colonisation and settlement of many island plant lineages remain elusive, our results are consistent with palaeodrought as a potential driver for woodiness in most of the insular woody Canary Island lineages studied.

## 1. INTRODUCTION

Since Darwin’s work on the Galapagos (1859), marine islands have been regarded as ideal systems to unravel evolutionary processes that have shaped present’s day life due to their isolation and defined boundaries (Losos & Ricklefs, 2009; Helmus et al., 2014; Patiño et al., 2017). Despite comprising just 3.5% of the Earth’s land area, marine islands – including volcanic islands, continental fragments, and land-bridge islands – harbour up to 20% of all terrestrial species and thereby contribute disproportionately to global biodiversity. This implies that a large proportion of island species evolved through speciation *in-situ* and are found nowhere else in the world, highlighting marine islands as natural laboratories of evolution (Whittaker et al., 2017). A famous example of the particularity of island biota is the repeated evolution of a peculiar suite of convergent traits in insular clades, often called “island syndromes”; examples are tendency toward flightlessness in birds and insects and loss of dispersal powers in plants, naïve behaviour and relaxed defences toward predators, body size changes, and insular woodiness (Burns, 2019).

Within flowering plants, woodiness is considered to be ancestral (Field et al., 2004; Doyle, 2012), meaning that herbaceous lineages lost woodiness that characterized their ancestrally woody ancestors. Surprisingly, many herbaceous lineages have evolved back into woody shrubs or even trees on islands as well as continents, an evolutionary reversal referred to as (phylogenetically) derived woodiness (Carlquist, 1974; Givnish, 1998; Lens, Davin et al., 2013). Insular woodiness in a strict sense – defined as the evolution of the woody growth form on islands after colonisation of herbaceous plant colonisers – is the best known type of derived woodiness. The remarkable prevalence of insular woodiness across (sub-)tropical islands helps to explain why these islands are proportionally woodier than nearby continents, which is without doubt the most striking feature of these insular floras (Carlquist, 1974; Whittaker & Fernández-Palacios, 2007; Lens, Davin et al., 2013). This suggests that (sub-)tropical islands are exceptionally suited to boost wood formation in otherwise herbaceous lineages, which has been baffling scientists ever since the original observations of Charles Darwin (Darwin, 1859) and Joseph Dalton Hooker (Hooker, 1866). However, information on derived woodiness across the world remains very fragmented. Therefore, the corresponding author has compiled a database including all known derived woody species in flowering plants based on rigorous screening of the life form traits (woody vs herbaceous) via taxonomic revisions and floras for species that have been included in hundreds of published molecular phylogenies. This huge effort resulted in a comprehensive global derived woodiness database revealing that evolutionary transitions from herbaceousness towards derived woodiness on continents have occurred more frequently than on islands (Lens & Zizka, unpublished dataset).

The causes of rampant convergent evolutionary transitions towards insular woodiness across the world have been debated for over 150 years (Whittaker & Fernández-Palacios, 2007; Lens, Davin et al., 2013). Why herbaceous colonisers develop into shrubs or even trees on (sub)tropical islands has been identified as one of the major questions that remain at the forefront of island studies 50 years after the ground-breaking publication of MacArthur and Wilson’s *Theory of Island Biogeography* (Patiño et al., 2017). A number of hypotheses have been put forward to explain insular woodiness, such as (i) increased competition (Darwin, 1859, elaborated by Givnish, 1998: taxon-cycling hypothesis), (ii) greater longevity (Wallace, 1878; elaborated by Böhle et al., 1996: promotion-of-outcrossing hypothesis), (iii) favourable climate (especially lack of frost; Carlquist 1974), and (iv) reduced herbivory (Carlquist 1974; for a more detailed explanation of these hypotheses, see Whittaker & Fernández-Palacios, 2007). However, experimental tests for these hypotheses are scarce and based on only a few, small-scale examples (Percy & Cronk, 1997; Givnish, 1982).

More recently, Lens, Davin et al. (2013) proposed a fifth hypothesis postulating drought as a potential driver for the evolution of wood formation, following two main lines of evidence. Firstly, the global derived woodiness database shows that most derived woody species (on continents and islands) are native to vegetation types with a marked drought season, such as Mediterranean regions, steppes and savannas, and (semi-)deserts (∼75% of the 7000 derived woody species - Lens & Zizka, unpublished dataset). Secondly, increased resistance of stems to drought-induced embolism inside water conducting cells, a trait that is associated with drier habitats and survival during drought (Lens, Tixier et al., 2013; Anderegg et al., 2016; Choat et al., 2018), seems correlated to increased woodiness. This is based on water transport measurements in stems comparing herbs and their derived woody relatives in *Arabidopsis* (Lens, Tixier et al., 2013) and Canary Island Asteraceae (Dória et al., 2018), as well as positive correlations between embolism stress resistance and stem lignification in herbaceous Canary Island Brassicaceae (Dória et al., 2019) and across grasses (Lens et al., 2016).

A detailed insular woodiness review paper based on molecular phylogenetic insights is only available for the Canary Islands, a volcanic archipelago about 100km off the coast of northwest Africa (Lens, Davin et al., 2013). Two important conclusions can be drawn from this review: (1) at least 220 native insular woody species are the result of 38 independent origins towards derived woodiness on the Canaries (belonging to 34 genera), and (2) the majority of the insular woody species typically grow in the markedly dry coastal regions, agreeing with the drought stress hypothesis. However, the link between insular woodiness and drought on the Canary Islands based on current-day distribution patterns does not necessarily mean that woodiness originated in drier palaeohabitats.

The main objective of this paper is to investigate whether the origin of insular woodiness amongst multiple independent island lineages is consistent with palaeoclimatic aridification episodes for the Canary Islands. Therefore, we reviewed the palaeoclimatic history for the archipelago based on the literature, and we used the largest dated angiosperm phylogenetic framework (Janssens et al., 2020) to simultaneously date colonisations and origins of woodiness in 24 (out of in total 38) insular woody lineages in order to place these insular woodiness transitions in the context of palaeoclimate. In addition, we provided dating estimates for the colonisation events of 13 additional Canary Island lineages representing nine herbaceous lineages that remained herbaceous on the archipelago, two ancestrally woody lineages, and two woody lineages that developed their woodiness outside the Canary Islands. As a secondary objective, the dating estimates of the 37 Canary Island lineages investigated were also linked to the number of native species on the archipelago in order to test the niche pre-emption hypothesis, stating that older lineages should be more species rich because they have had more opportunity to occupy the available niches compared to younger lineages (Silvertown 2004; Silvertown et al., 2005).

## 2. MATERIALS AND METHODS

### 2.1 Marker choice and taxon sampling

The Consortium for the Barcode of Life working group (CBOL) recommended the two chloroplast markers *matK* and *rbcL* as primary barcodes to identify plant species, resulting in massive amounts of sequences available in GenBank. The combination of both markers maximizes phylogenetic resolution in a complementary way, since *rbcL* is a conservative locus and therefore useful for reconstructing deeper nodes in phylogenies, while *matK* contains rapidly evolving regions that are useful for resolving shallower nodes (Hollingsworth et al., 2011; Janssens et al., 2020). Moreover, working with only two markers also reduces missing data in large datasets to a minimum, as it is known that missing data could lead to wrongly inferred relationships (Roure et al., 2013). Downside of this approach is, however, that the resulting *rbcL-matK* gene tree is not necessarily the same as the corresponding species tree, due to processes such as hybridization, incomplete lineage sorting and horizontal gene transfer (Davidson et al., 2015). For this reason, we only worked with Canary Island lineages for which the phylogenetic relationships matched those in previously published phylogenies based on various plastid-nuclear genes (see below).

Many of the available *rbcL-matK* sequences in GenBank were mined to compile a large-scale angiosperm dataset including 36,101 species, to which 56 fossils were assigned as calibration points (Janssens et al., 2020). This dataset allowed us to have an evolutionary framework to estimate the colonization times of the Canary Island lineages using a single analysis (see below). We modified this original dataset to meet our research question, a focus on the eudicots since most insular woody transitions on the Canaries belong to the asterid and rosid clades, and we excluded nearly all monocots from our sampling because these species never produce wood, making them redundant for our purpose. This reduced our dataset to 23,781 species associated to 42 fossil calibration points. We included sequences of *matK* and *rbcL* from an additional 135 Canary Island species that are endemic, native or probable native according to Arechavaleta et al. (2010). Of these species, 95 are insular woody according to the review paper by Lens, Davin et al. (2013). We also included sequences of both markers for 103 continental outgroup species (see Appendix S1 in Supporting Information) that were found to be the most closely related to the native Canary lineages based on their position in previously published phylogenies. Of these 238 species in total, we generated original *matK* and *rbcL* sequences for 100 Canary Island species and 24 continental outgroup species, collected from dried leaf samples or DNA samples via different institutes: (1) DNA Bank of the Canarian Flora and LPA herbarium, housed at the Botanical Garden Viera y Clavijo - Unidad Asociada al CSIC, Cabildo de Gran Canaria (Canary Islands, Spain), (2) herbaria of Naturalis Biodiversity Center (Leiden, The Netherlands), Meise Botanic Garden (Belgium), La Laguna University (Tenerife, Canary Islands, Spain), and Texas (USA), and (3) botanical gardens of Leiden (The Netherlands), Utrecht (The Netherlands), and Marburg (Germany). Additionally, marker sequences of 35 Canary Island species and 79 continental outgroup species were downloaded from GenBank. Vouchers used in our study are compiled in Appendix S1 in Supporting Information.

Despite our strategy for outgroup sampling and the expected distribution range for most of the outgroup species in the Mediterranean region (Appendix S1 in Supporting Information), it is likely that for several Canary Island lineages the closest relatives have not yet been described or went extinct. Therefore, the age estimates for island colonisation should be considered as maximum ages and may overestimate the true colonisation time, especially when stem ages are used instead of crown ages. We opt to mainly base our discussion on stem age estimates (Fig. 1), but also integrate estimates from crown node ages (Fig. S1 in Supporting Information) to have a more balanced discussion on colonisation of island lineages (for a more detailed discussion, see García-Verdugo et al., 2019).

**Figure 1:**
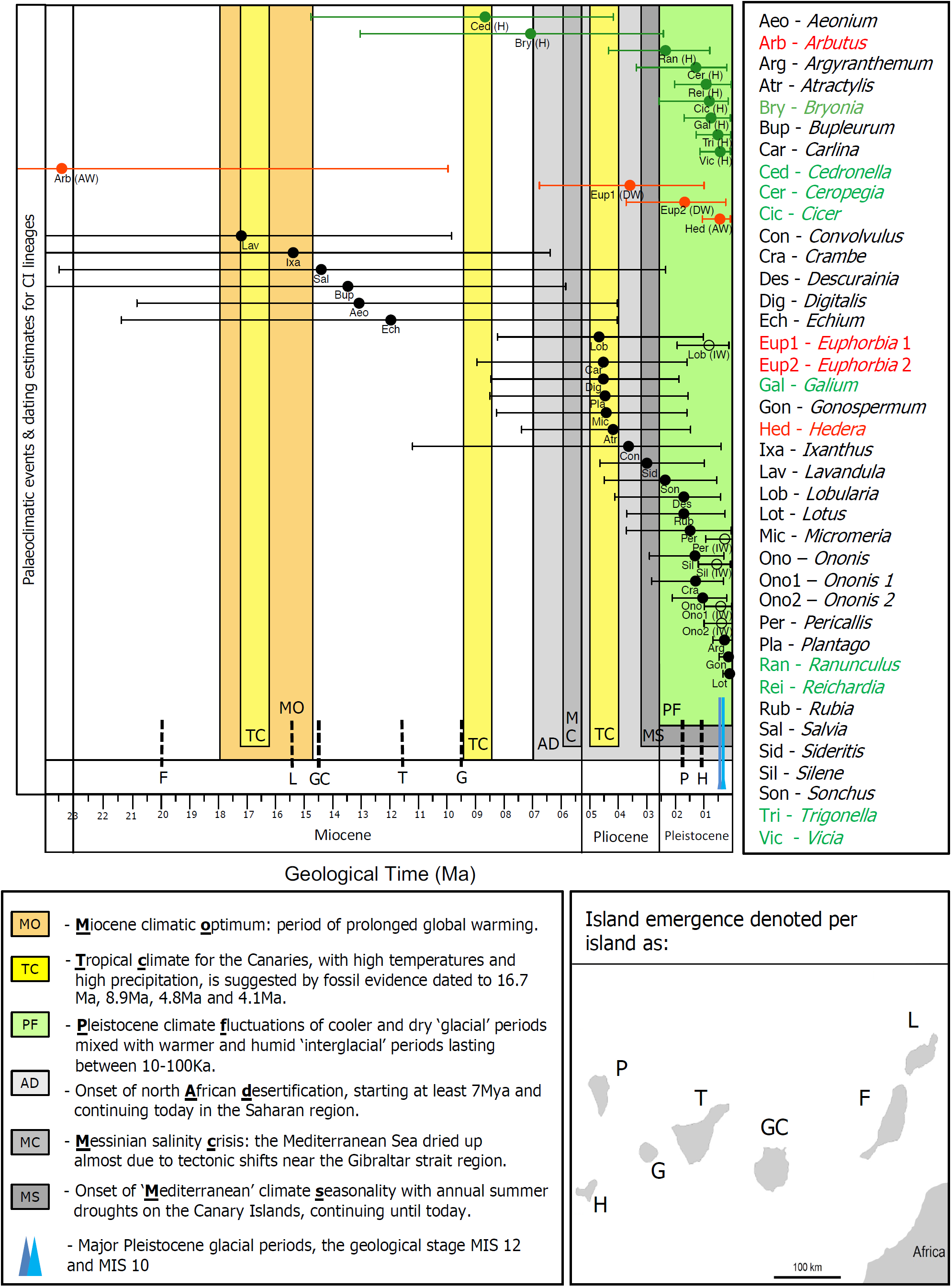
Timeline of 37 Canary Island colonisation events and insular woody shifts based on conservative mean stem age estimates and their 95% range, along with the major palaeoclimatic events, and ages of the individual Canary Islands over the last 23 Myr. Black circles refer to the estimated mean ages for the timing of colonisation of the insular woody clades; for *Lobularia, Ononis, Pericallis* and *Silene*, which all include herbaceous as well as insular woody species on the Canary Islands, the insular woody shift occurred more recently in time (open circles). The time of colonisation of the herbaceous clades are illustrated in green and denoted by ‘H’, and the time of colonisation of ancestrally woody (‘AW’) clades and derived woody (‘DW’) clades that have evolved their woodiness outside the archipelago are illustrated in red.

In total, the species investigated belong to 37 independent, single colonisation events and 18 flowering plant families: 24 insular woody lineages (63% of the total number of insular woody lineages on the Canaries), 9 herbaceous eudicot lineages, 2 derived woody lineages (developed their woodiness outside the Canaries), and 2 ancestrally woody lineages. For most of the nine native herbaceous Canary Island lineages, we selected clades from the same families that also included the insular woody lineages. We did not retain a number of additional clades for which we generated sequences, because the *matK-rbcL* phylogeny did not match the previously published phylogenies based on various plastid or nuclear markers. For instance, *matK-rbcL* sequences did not discriminate the Canary Island clade with the continental outgroup (e.g. *Canarina, Cheirolophus, Rumex, Urtica*), or the lineage was not found to be monophyletic in the *rbcL-matK* consensus topology as opposed to previously published phylogenies. For the *Aeonium* alliance, we could only provide the dating estimate for the colonisation event, because the phylogenetic position of the insular woody *Aeonium* species did not match the well-supported position found in previous phylogenies.

### 2.2 DNA extraction, sequencing protocols and alignment

For each specimen a 1cm^2^ subsample of leaf tissue was obtained. The leaf material was lysed with sterile sand and 7mm glass beads before DNA was extracted with the NucleoMag 96 plant kit (Macherey-Nagel Gmbh & Co., Düren, Germany) using the KingFisher Flex magnetic particle processor (Thermo Scientific). The primer pair 1R-KIM/3F-KIM (forward and reverse) was used for *matK* amplification (CBOL Plant Working group, 2009). For *rbcL*, a 900bp sequence was amplified with the forward primer “F1” and an internal reverse primer; another 800 bp sequence was amplified with an internal forward primer and the *rbcL* “85R” or “Z1375R” reverse primer (Kress & Erickson, 2007). Both DNA regions were amplified using an optimised PCR mixture consisting of 8.4 µl ultrapure H_2_O, 5.0 µl 5× PCR Phire reaction buffer, 1.0 µl 25 mM MgCl2, 1.0 µl 10mg/ml BSA, 5 µl 100 mg/ml PVP, 0.5 µl 2.5 mM dNTPs, and 0.5 µl of Phire Hot Start II polymerase. A similar hot start PCR protocol was used for *matK* and *rbcL*, only differing in the annealing temperature (53 °C and 66 °C, respectively). Thermal cycling conditions were as follows: 98 °C for 45 seconds (s), followed by 40 cycles of 98 °C for 10 s, 53 °C or 66 °C for 30 s, 72 °C for 40 s, and a final extension at 72 °C for 7 min. The PCR products were verified by electrophoresis in 2% agarose e-gel stained with SYBR™ Safe DNA Gel Stain (Thermo Scientific).

The PCR products were purified and sent to BaseClear (Leiden, The Netherlands) for bi-directional Sanger sequencing. Consensus sequences were generated and edited manually using Geneious version 11.1 (https://www.geneious.com, Kearse et al., 2012). The concatenated *matK* and *rbcL* sequences were added into an existing, aligned angiosperm-wide dataset (Janssens et al., 2020). To facilitate the aligning process of our self-generated sequences to the sequences already aligned in the dataset, we aligned each obtained sequence of a particular clade to a selection of phylogenetically related reference sequences in the existing dataset using MAFFT version 7 (with parameters keep alignment length: yes, progressive method: G-INS-1, all other parameters default) (Katoh & Standley, 2013). Manual adjustments were made using Geneious.

### 2.3 Construction of a dated maximum likelihood phylogenetic tree

We used the CIPRES scientific portal to run RAxML - HPC version 8 (Miller et al., 2010; Stamatakis, 2014, 2016), GTRCAT approximation was selected in accordance with RAxML recommendations for large trees following Izquierdo-Carrrasco et al. (2011), the other default parameters followed Stamatakis (2016). The phylogenetic relationships of all species in the broad-scale aligned dataset were constrained at the family level when conducting the RaxML analysis, in order to (1) assure that the discriminatory power of *matK* and *rbcL* at the genus and species level comes at the cost of correct deeper phylogenetic reconstruction, and (2) because of reduced computational time.

Fourty-two age constraints across the angiosperm phylogeny were taken from Janssens et al (2020; for phylogenetic positions, see Appendix S2 in Supporting Information). The root node was calibrated to 138.5 Mya (million years ago) based upon recent estimates by Magallón et al. (2015). Molecular dating was completed using TreePL version 1.0, which is specifically designed to date large phylogenies (Smith & O’Meara, 2012). A small smoothing parameter of 0.003 was applied to minimise the influence of the large heterogeneity of substitution rates across the broad angiosperm phylogeny, while the default setting was kept for all other parameters (for more details to justify the smoothing parameter, see Janssens et al., 2020). Changing the smoothing parameter to 1 or even 10 did not dramatically change the dating estimates for most of the clades studied (results not shown).

To assess the phylogenetic uncertainty in our dataset, 200 trees were created using RAxML – each with a unique starting position in the “hill climbing” space – and independently dated using the same set of age priors with TreePL. The 200 phylogenetic trees were summarised into a single consensus tree that was used to calculate the 95% range for the dating estimates of each node using TreeAnnotator version 1.8.4 with default settings (Drummond et al., 2012). The consensus tree summarizing the 200 maximum likelihood trees is given in annotated nexus format in Appendix S3 in Supporting Information and available on Dryad (https://doi.org/10.5061/dryad.kh189322s).

An ancestral state reconstruction for our 24,000 species tree was not performed, because it is important to have a broad (herbaceous) outgroup sampling for each Canary Island lineage studied. Since we only targeted the closest relatives known based on previous phylogenies to estimate colonization times, an ancestral state reconstruction of the angiosperm wide tree could be problematic for at least some of the insular woody lineages studied that clearly underwent an evolutionary transition from herbaceousness towards woodiness. Therefore, we relied on the information provided in the insular woodiness review paper by Lens, Davin et al. (2013) and the updated global derived woodiness database (Lens & Zizka, unpublished dataset), and presented means and confidence ranges of stem and crown node estimates for 24 insular woody shifts and their colonisation events (of which 19 are equal to the estimated date of the insular woody shift since all island species are insular woody; Table S1 in Supporting Information). In addition, we added similar information for nine herbaceous, two ancestrally woody and two derived woody Canary lineages that acquired their (derived) woodiness before colonisation on the archipelago (Table S1), and visualised the mean stem age estimates and their confidence ranges with respect to the palaeoclimatic timeline in Fig. 1. All 37 Canary Island clades investigated were supported by support values greater than 0.75 in the consensus tree based on 200 replications with unique starting position, with a majority of clades having support values greater than 0.95. The dates for the crown and stem node lineages were compared to dates from the literature when available (Table S1).

To test the niche pre-emption hypothesis, we plotted the number of native Canary Island species to the estimated colonization time for all 37 Canary Island clades and applied a linear regression analysis using base statistical and graphical functions of R (v3.5.1., R core team, 2018) (Fig. 2). Species numbers are derived from the Canary Island species list (Arechavaleta et al., 2010), unless otherwise stated.

**Figure 2:**
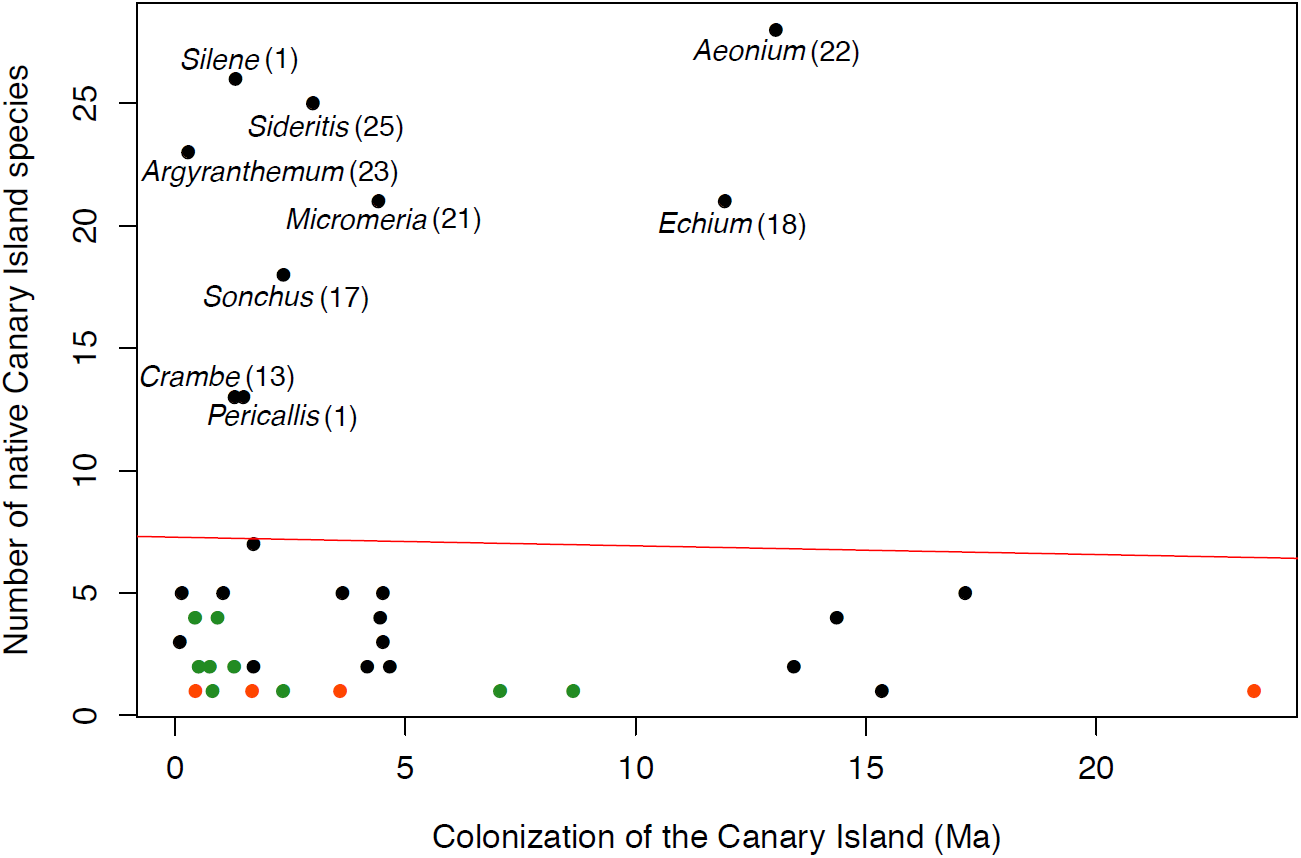
Graph showing no relationship between mean stem age estimates of the colonisation of the 37 Canary Island clades and their species richness based on the simultaneous dating analysis. Linear regression is indicated by the red line (slope = −0.035, R^2^ = 0.00051). Data points are coloured according to growth form: clades with at least one insular woody species (black), entirely herbaceous lineages (green), derived woody lineages developed woodiness outside Canaries or ancestrally woody lineages (red). Data points of the largest clades are labelled and the number of native insular woody species in these large clades is indicated in brackets.

### 2.4 Palaeoclimatic reconstruction

A timeline of global, Mediterranean/North-African, and Canary Island-specific palaeoclimatic trends and events was compiled after a review of published literature (Appendix S4 in Supporting Information). Diverse sources of evidence were considered to reconstruct the palaeoclimate for the Canaries, including fossilised pollen, oceanic benthic foraminifera cores, results from geochemical or geological analyses, faunal trace fossils and atmospheric models. Highlights of this elaborate dataset were visualized in Fig. 1.

## 3. RESULTS

### 3.1 Molecular dating estimates

The colonisation of the Canary Islands is inferred to be relatively recent for the clades studied, even with the most conservative dating estimates based on stem ages, with most events occurring in the last 3.2 million years (Myr) for both herbaceous and ancestrally woody lineages (7/9 and 2/4 lineages for stem age estimates, respectively; Fig. 1). With respect to dating events of the shifts towards insular woodiness, 19 of the 24 transitions studied have likely occurred during the last 7 Myr (80%), taking into account the large confidence intervals of the mean stem ages. Out of these 24 insular woody shifts, 13 (55%) are estimated to have occurred within the last 3.2 Myr. Using crown node estimates (Table S1 and Fig. S1 in Supporting Information), 80% of the ancestrally woody lineages would have originated during the last 3.2 MY, 21 of the insular woody transitions would have occurred within the last 7 MY (ca. 90%), 16 (65%) of these during the last 3.2 MY.

### 3.2 Palaeoclimate review for the Canary Islands

A literature overview of the major palaeoclimatic events during the last 20 Myr for the Canary Islands, North Africa and Mediterranean region, and a few global trends are presented in Appendix S4 in Supporting Information. The most important events that have impacted the palaeoclimate on the Canary Islands are visually summarised in Fig. 1, which also illustrates long periods of uncertainty about the Canarian palaeoclimate. The two major palaeoclimatic events which have caused severe aridification episodes on the Canaries include (i) the Pleistocene fluctuations with drier glacial cycles of which the two main cycles date back 0.38 – 0.34 million years ago (Mya) and 0.48 – 0.42 Mya, and (ii) the onset of a Mediterranean seasonality on the Canaries with annual warm dry summers and mild wet winters (3.2 Mya). Furthermore, the Messinian salinity crisis leading to a massive drying event of the Mediterranean Sea (5.96 – 5.33 Mya), and the onset of desertification of Northern Africa (∼ 7 Mya) have likely impacted the Canary Island vegetation.

### 3.3 Age and species richness of insular woody clades

We found no relationship (not based on stem nor crown node age) between the divergence time estimates of the 37 Canary Island lineages and their current species richness on the Canaries based on Arechavaleta et al. (2010) (linear regression; slope = −0.035, R^2^: < 0.001; Fig. 2). For instance, there are a number of recent, rapid radiations among some of the largest insular woody Canary Island clades: *Argyranthemum* includes 23 insular woody species and has radiated 0.29 Mya (CI: 0.69 – 0.01 Mya), *Silene* has radiated into 26 species (of which eight endemic, two likely native, and 16 possible native with much uncertainty about their origin) in the last 1.32 Mya (CI: 2.91 – 0.31 Mya), *Sonchus* includes 17 insular woody species and has originated 2.36 Mya (CI: 4.49 – 0.56 Mya). Additionally, the three oldest clades recovered from our molecular dating analysis reveal three relict lineages: the monospecific insular woody *Ixanthus* dates back 15.35 Mya (CI: 24.53 – 6.37 Mya), the insular woody clade of *Lavandula* dates back 17.16 Mya (CI: 26.96 – 9.82 Mya), and the sole native, ancestrally woody *Arbutus* (*A. canariensis*) has an estimated age of 23.43 Mya (CI: 42.30 – 9.94 Mya).

## 4. DISCUSSION

To our knowledge, this work presents the largest dating study for island plants so far. We provide comparable dating estimates for the colonisation of 37 Canary Island lineages, including 24 (out of in total 38) insular woody lineages native to the Canaries (Lens, Davin et al., 2013), based on the recent megaphylogeny for flowering plants (Janssens et al., 2020) that makes use of two chloroplast markers and 42 fossil calibration points outside the Canaries. Even considering that our age estimates are upper bounds based on mean stem ages, our analysis highlights the young age of most Canary Island lineages studied, dating back to the Plio-Pleistocene (Fig. 1). For 70% of the lineages studied, the origins of these extant Canary Island lineages are in line with previously published island studies using alternative calibration points (e.g., different fossils, age of islands, or different secondary calibration points) and different nuclear and or chloroplast markers (Kondraskov et al., 2015; García-Verdugo et al., 2019), making our large-scale dating approach valid (Table S1). However, we are aware that the use of only two chloroplast markers for phylogenetic reconstruction could lead to erroneous species relationships. Therefore, we only selected the clades of interest from the Canaries in the large-scale angiosperm phylogeny that presented correct outgroup-ingroup relationships, as defined by previously based phylogenies based on various chloroplast and/or nuclear markers. Until a more complete phylogeny is available for flowering plants based on a higher genomic coverage, the proposed two-gene phylogenetic framework (Janssens et al., 2020) represents a useful tool that has already been successfully applied to answer broad-scale evolutionary questions across angiosperms (Dagallier et al., 2020).

To assess whether the woodiness transitions across the 24 Canary Island insular woody clades studied are consistent with palaeoclimatic aridification periods, we matched the age estimates with a palaeoclimatic review of the Canaries that we compiled from the literature (Fig. 1; Appendix S4 in Supporting Information). When tracing trait shifts or lineages back to the palaeoclimate in which they evolved, two sources of uncertainty must be accounted for. Firstly, the estimated ages obtained for the insular woody clades are accompanied by large confidence intervals, which could make the link to a specific palaeoclimatic episode blurry (Fig. 1). One way to extract a general pattern from these estimates is to investigate multiple lineages, as we did here. A second point of concern is that it is hardly possible to find out in which microhabitat the first coloniser(s) settled on a specific island. Especially for high elevation islands, there is a large altitudinal range in vegetation types ranging from dry coastal scrub vegetation resembling (semi-)deserts to higher-elevation laurel forests that receive daily precipitation from clouds carried by the north-eastern trade winds (del Arco et al., 2006). Furthermore, the archipelago’s geo-ecological complexity has changed dramatically through time, and there were high elevation islands in the past that lost their relief and many habitats due to collapses and landslides (Caujapé-Castells et al., 2017). In this challenging context, one way to estimate the type of original colonised habitat is to perform ancestral area reconstructions based on phylogenies of current-day species. In this regard, it is interesting to see that the dry lowland scrub was found to be the ancestral Canary habitat for the insular woody genera including *Crambe* (Francisco-Ortega et al., 2002), *Descurainia* (Goodson et al., 2006), and *Echium* (García-Maroto et al., 2009).

### 4.1 Repeated evolutionary transitions towards derived woodiness are recent and consistent with palaeoclimatic aridification events on the Canary Islands

While there is evidence for at least four insular woody relict lineages dating back to more than 13 Mya (*Ixanthus, Bupleurum, Lavandula* and *Salvia*), our conservative dating results based on stem age estimates indicate that the evolution towards woodiness in 80% of the insular woody clades studied likely originated during the last 7 Myr (Fig. 1). When crown ages are taken into account, up to ca. 90% (19 out of 24) of the insular woody lineages probably evolved within the last 7 Myr (Fig. S1 in Supporting Information). In other words, most native Canary Island lineages prove to be relatively young, which is confirmed by the review study of García-Verdugo et al. (2019) who found that the mean crown node ages of Canarian plant lineages date back 2.1 ± 2.4 Mya, with most of the repeated transitions towards insular woodiness evolving from the late Miocene (stem age estimates) or early Pliocene (crown age estimates) onwards. The late Miocene is characterized by two major aridification events that happened in close proximity to the Canaries: (1) the onset of desertification of Northern Africa (∼ 7 Mya) and (2) the Messinian salinity crisis leading to a massive drying event of the Mediterranean Sea due to closure of the Strait of Gibraltar (5.96 – 5.33 Mya). These two aridification events outside the Canaries have likely impacted the vegetation on the archipelago, because of both its proximity to the mainland and the prevailing north-eastern trade winds that help explain the Mediterranean origin for most of the native Canary Island lineages. Moreover, the origin of woodiness in 55-65% of the insular woody clades studied originated during the late Pliocene and Pleistocene, a palaeoclimatic timeframe that coincided with two additional aridification events characteristic for the Canaries: (1) the onset of the Mediterranean climate with annual warm summer droughts starting 3.2 Mya, likely caused by the redirection of oceanic currents of the Atlantic Ocean after the closing of the Panamanian Isthmus (Haywood et al., 2000; Jiménez-Moreno et al., 2010; Meco et al., 2015), and (2) the drier glacial periods during the Pleistocene, in particular two glacial maxima during 0.48 – 0.42 Mya and 0.38 – 0.34 Mya (see next paragraph; Fig. 1).

The two short glaciation events that occurred in quick succession of each other between 0.48 and 0.34 Mya are considered to be the most severe glacial periods across the Northern Hemisphere over the last 2 million years, and are defined by the marine isotope stages (MIS) 12 and 10, respectively (Shackleton, 1987; Lisiecki & Raymo, 2005; Sánchez Goñi et al., 2016). The mean colonisation estimate of the insular woody *Argyranthemum* based on stem age (0.29 Mya) is consistent with MIS 10, which may imply that this relatively short but intense aridification episode could have driven woodiness and subsequent fast radiation in this genus leading to 23 Canary Island species (and four additional species on Madeira and the Selvagens Islands). This suggested link also matches the experimental study by Dória et al. (2018) showing that the stems of the woody *Argyranthemum* species are better able to avoid drought-induced air bubble formation inside water conducting vessels compared to stems of herbaceous continental relatives. Even if *Argyranthemum* was older (1.5-3.0 Mya based on divergence times from isozymes and from chloroplast restriction site DNA and a different outgroup sampling; Francisco-Ortega et al., 1995, 1996, 1997), the same link between insular woodiness and palaeodrought could be made. Additionally, the estimated timing of insular woody shifts in *Gonospermum* (CI: 0.48 – 0 Mya), *Pericallis* (CI: 0.94 – 0.01 Mya), two shifts in the *Ononis* Canary Island clade (CI: 1 – 0.02 Mya; 0.99 – 0.04 Mya), *Silene* (CI: 1.2 – 0.08 Mya) and *Lobularia* (CI: 1.94 – 0.13 Mya) may have overlapped the two marked glacial cycles. On the other hand, the colonisation of the herbaceous Canary Island lineages *Vicia* (CI: 1.14 – 0.08 Mya), *Trigonella* (CI: 1.28 – 0.07 Mya), *Galium* (CI: 1.7 – 0.08 Mya), *Cicer* (CI: 2.57 – 0.16 Mya), *Reichardia* (CI: 2.03 – 0.05 Mya) and *Ceropegia* (CI: 3.36 – 0.21 Mya) could also be attributed to MIS 10 or MIS 12, but we do not know whether these herbaceous lineages colonised and initially diversified in the same micro-niches as the insular woody clades with comparable age.

Some of the insular woody clades have originated very recently, such as *Lotus* (0.11 Mya) and *Gonospermum* (0.15 Mya), suggesting that the transition from the herbaceous towards the woody growth form can evolve very rapidly. One of the most intriguing examples of this rapid evolution towards woodiness can be found on the Canary Islands, where humans introduced *Brassica oleracea* (cabbage), leading to the spectacular woody, ‘walking stick’ accession (Lens, Davin et al., 2013). The exact timing of human introduced cabbage remains unclear: there is no archaeological evidence of the presence of cabbage in the aboriginal time from 3^rd^ century BC to 15^th^ century AC (Morales et al., 2009, 2017), although there is one reference quoting orchards of cabbages cultivated on the islands in 1341 AC (Barker-Webb & Berthelot, 1842). This means that cabbages could have been introduced roughly 700 years ago at the end of the aboriginal time via trade with the Florentines, Genoese, Spaniards and Portuguese. Therefore, it is not surprising that a number of clades have undergone major evolutionary changes on the Canaries in recent geological times.

### 4.2 The remarkable link between insular woodiness and diversification on the Canaries

Out of the ten largest angiosperm genera of the Canary Islands [viz. *Aeonium* (28 sp.), *Sideritis* (25 sp.), *Limonium* (22 sp.), *Echium* (21 sp.), *Micromeria* (21 sp.), *Argyranthemum* (23 sp.), *Cheirolophus* (18 sp.), *Lotus* (19 sp.), *Sonchus* (18 sp.) and *Crambe* (13 sp.)], eight include several insular woody species accounting for 147 out of 220 insular woody species on the Canary Islands (67%), while the herbaceous genera mostly do not harbour more than five species (*Silene* is a noteworthy exception; Arechavaleta et al., 2010). Thus, *in-situ* wood development on the Canaries may have boosted species diversification, potentially via improved drought stress resistance compared to herbaceous species (Dória et al., 2018). However, this seems to contradict outcomes of large-scale comparative studies showing that herbaceous lineages have 3-4x as many species than (ancestrally) woody sister lineages, which likely reflects the shorter generation time of herbs often leading to significantly higher rates of molecular evolution compared to tall trees (Smith & Donoghue, 2008; Givnish, 2010). Since it is plausible that rates of molecular evolution between *insular* woody lineages (mostly comprised of small shrubs) and their herbaceous continental relatives (including many perennial herbs) resemble each other more, the observed link between diversification and insular woodiness on the Canaries may not be in conflict with the large-scale comparative studies described above.

Identifying the traits that promote diversification is one of the unresolved key questions in plant evolutionary sciences (Patiño et al., 2017; Sauquet & Magallon, 2018). The potential drivers behind species radiation are the result of complex ecological relationships, and probably involve a range of abiotic (soils, climate, geology) and/or biotic traits (flower, seed, fruit, ploidy) that may or may not have co-evolved with the insular woody life form and are perhaps more likely to cause geographic (and thus reproductive) isolation leading to speciation (Givnish, 2010; Caujapé-Castells et al., 2017). Likewise, drought is thought to have driven the diversification of ancestrally woody lineages as well, for example in gymnosperms (Pittermann et al., 2012; Larter et al., 2017), but it is important to add that not all woody species are drought-tolerant. For example, several of the ancestrally woody Canary Island lineages are native to the wet laurel forests (e.g., *Apollonias, Laurus, Morella, Ocotea, Persea*), and presumably sensitive to drought.

The insular woody clades in our dating analysis show a wide range of mean stem ages throughout evolutionary time: *Ixanthus* (15.35 Mya), *Echium* (11.94 Mya), *Micromeria* (4.42 Mya), *Sideritis* (3 Mya), *Sonchus* (2.36 Mya), *Crambe* (1.30 Mya), *Argyranthemum* (0.29 Mya). Surprisingly, these time estimates do not correlate with the number of species in each insular woody lineage, with some of the oldest lineages (*Bupleurum*, 2 sp.; *Ixanthus*, 1 sp.) having far fewer species than some of the recent lineages (*Argyranthemum*, 23 sp.; *Crambe*, 13 sp.; *Sideritis*, 25 sp.). Also, in non-insular woody lineages, there is no link between number of species and colonisation time: the herbaceous *Vicia* (0.44 Mya) has diversified into four native species, while the ancestrally woody *Arbutus* (23.43 Mya) has only one native species. In other words, there is no relationship between species richness and clade stem age across all the 37 Canary Island lineages studied (Fig. 2). These results are in conflict with the niche pre-emption hypothesis stating that early colonisers have had more opportunity to occupy the available niches compared to later colonisers, and therefore older lineages should have had more chance to diversify into larger clades (Silvertown 2004, Silvertown et al., 2005). It is tempting to assume based on the relatively young age of most Canary Island lineages that the native flora of the archipelago is characterized by a high taxon turnover that is probably facilitated by high extinction rates on islands and the close proximity of mainland source areas (García-Verdugo et al., 2019).

In conclusion, we find that the woodiness transitions in about 80-90% of the insular woody lineages studied coincides with a period of palaeoclimatic aridification nearby the Canary Islands (starting 7 Mya), followed by the onset of annual summer droughts on the Canaries 3.2 Mya (about 55-65% of insular woody lineages studied), and with periods of intense aridification during the Pleistocene 0.48-0.34 Mya (about 30-40% of insular woody lineages studied). While a causal link between drought and wood formation cannot be confirmed based solely on our data, our results are consistent with the hypothesis that most evolutionary transitions towards woodiness on the Canary Islands took place in dry palaeoclimatic time frames. This agrees with experimental data on native current-day Canary Island species (Dória et al., 2018, 2019), and it confirms global distribution patterns of continental derived woody species across the world that often thrive in vegetation types with a marked drought season (Lens & Zizka, unpublished dataset). We hypothesise that the increased fitness of derived woody species in drought conditions compared to that of herbaceous species may have triggered diversification in several of the insular woody lineages. The use of explicit palaeoclimate data from earth system models, and the use of explicit models of trait evolution for derived woodiness, including phylogenetic null models, are promising approaches for future research to refine the evolutionary perspective on the link between derived woodiness and drought once more data is available. With this in mind, it is safe to assume that drought is definitely not the only potential driver for wood formation across all islands or all derived woody lineages (Kidner et al., 2016). Woodiness is a complex trait, which can be triggered by various understudied mechanisms, as demonstrated by the wide range of habitats – ranging from extremely dry to extremely wet areas – in which derived woody species are currently growing.

## Data availability

All DNA sequences newly produced have been deposited on GenBank with accession numbers MN783748 – MN783978 (see Appendix S1 in supporting information, for exact accession numbers). The phylogeny that support the findings of this study will be openly available in the DRYAD Digital Repository. Hooft van Huysduynen et al. (2019). Multiple origins of insular woodiness on the Canary Islands are consistent with palaeoclimatic aridification (https://doi.org/10.5061/dryad.kh189322s).

## ACKNOWLEDGEMENTS

All the collection institutes are acknowledged for sharing research material (Botanical Garden Canario ‘Viera y Clavijo’ - Unidad Asociada al CSIC (DNA Bank and LPA herbarium), Botanical Garden Marburg-Karteiauszug, Naturalis Biodiversity Center, Botanical Garden Meise, Botanical Garden Utrecht, Herbarium La Laguna University, Hortus Botanicus Leiden, Texas Herbarium).

## BIOSKETCH

Alex Hooft van Huysduynen is broadly interested in biodiversity patterns. This work represents a component of his MSc work on palaeoclimatic conditions for the Canary Islands during which insular woody groups have evolved. His internship was supervised by Frederic Lens who leads a team working on the evolutionary transition from herbaceousness towards derived woodiness (https://www.naturalis.nl/en/frederic-lens).

## Author contributions

F.L. and S.J. conceived the ideas; A.H.H., S.J., B.K., D.M., R.J.M., J.C.C., A.M.R., M.A. and F.L. collected the data; A.H.H., V.M., R.V., L.V., A.Z., M.L., J.M.F.P., L.N and F.L. analysed the data; and A.H.H. and F.L. led the writing, with contributions from everyone.

## Notes

### Competing Interest Statement

The authors have declared no competing interest.

