## Supporting information captions for "Multiple origins of insular woodiness on the Canary Islands are consistent with palaeoclimatic aridification"

**Appendix S1.** List of species for which matK and rbcL sequences are generated, along with their growth form, voucher information, GenBank number and collection institute.

**Appendix S2.** List of calibration points and references used for dating the angiosperm wide phylogeny.

**Appendix S3.** Annotated tree file in nexus format showing the consensus tree based on the

200 ML runs.

**Appendix S4.** Literature overview of palaeoclimatic events for the Canary Islands, northern Africa and the Mediterranean, and some global events during the archipelago formation.

**Table S1:** Summary of information for each Canary Island clade analysed. Mean molecular dating estimates and 95% confidence intervals of colonisation of the Canary clades and insular woody shifts are shown. Support values for phylogenetic topology from 200 maximum likelihood replications are also given. Also, the number of Canary Island and insular woody species in each lineage is indicated, as well as dating estimates from previously published phylogenies.

**Figure S1:** Timeline of 38 Canary Island colonisation events and insular woody shifts based on conservative mean crown age estimates and their 95% range, along with the major palaeoclimatic events, and ages of the individual Canary Islands over the last 23 MY. Black circles refer to the estimated mean ages for the timing of colonisation of the insular woody clades; for *Lobularia*, *Ononis*, *Pericallis* and *Silene*, which all include herbaceous as well as insular woody species on the Canary Islands, the insular woody shift happened more recently in time (open circles). The time of colonisation of the herbaceous clades are illustrated in green and denoted by ‘H’, and the time of colonisation of ancestrally woody (‘AW’) clades and derived woody (‘DW’) clades that have evolved their woodiness outside the archipelago are illustrated in red.
