## Supplementary material for "Multiple origins of insular woodiness on the Canary Islands are consistent with palaeoclimatic aridification": Figure S1

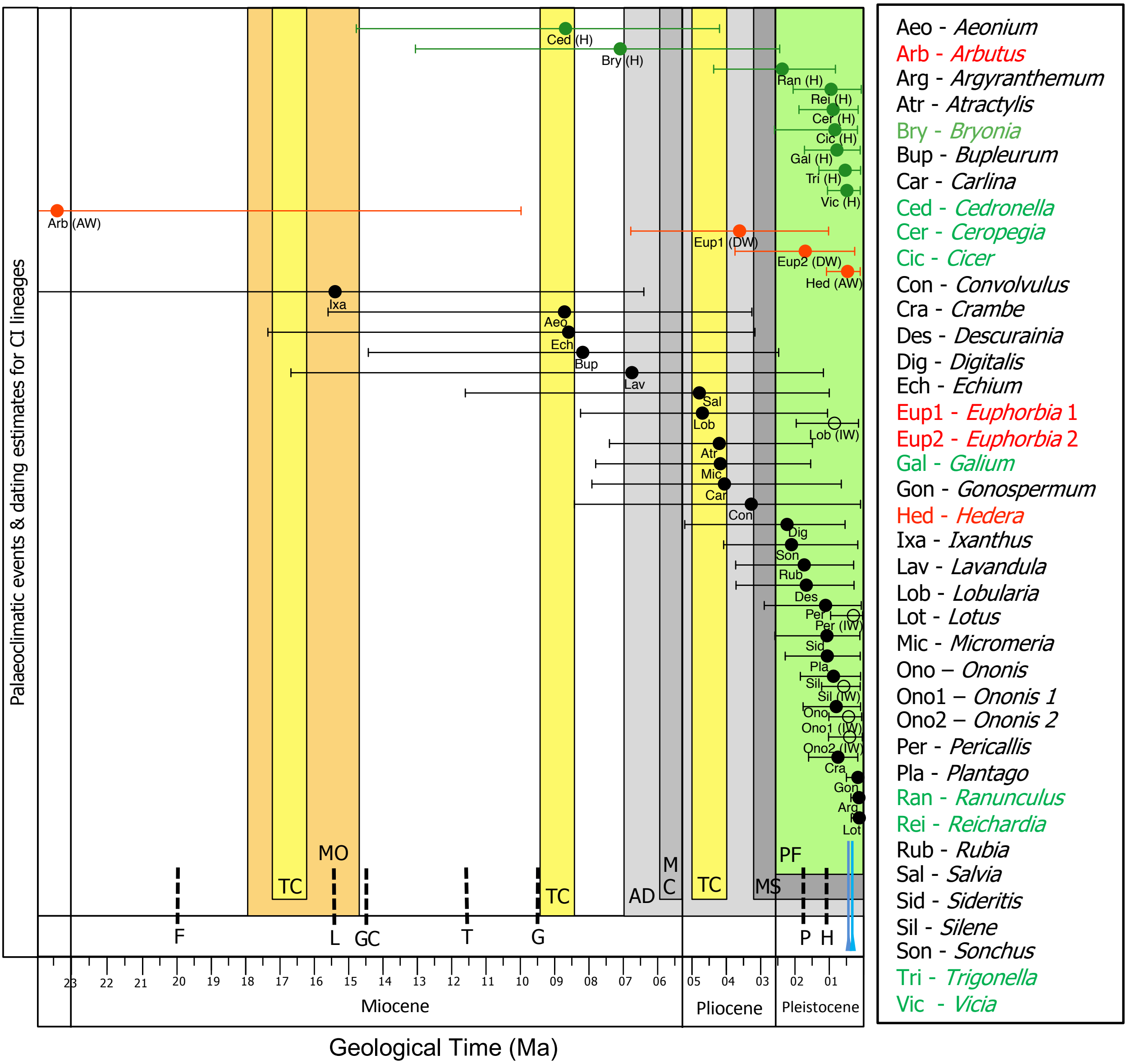


MO - **M**iocene climatic **o**ptimum: period of prolonged global warming.

TC - **T**ropical **c**limate for the Canaries, with high temperatures and high precipitation, is suggested by fossil evidence dated to 16.7 Ma, 8.9Ma, 4.8Ma and 4.1Ma.

PF - **P**leistocene climate **f**luctuations of cooler and dry 'glacial' periods mixed with warmer and humid 'interglacial' periods lasting between 10-100Ka.

AD - Onset of north **A**frican **d**esertification, starting at least 7Mya and continuing today in the Saharan region.

MC - **M**essinian salinity **c**risis: the Mediterranean Sea dried up almost due to tectonic shifts near the Gibraltar strait region.

MS - Onset of '**M**editerranean' climate **s**easonality with annual summer droughts on the Canary Islands, continuing until today.

- Major Pleistocene glacial periods, the geological stage MIS 12 and MIS 10

Island emergence denoted per island as:

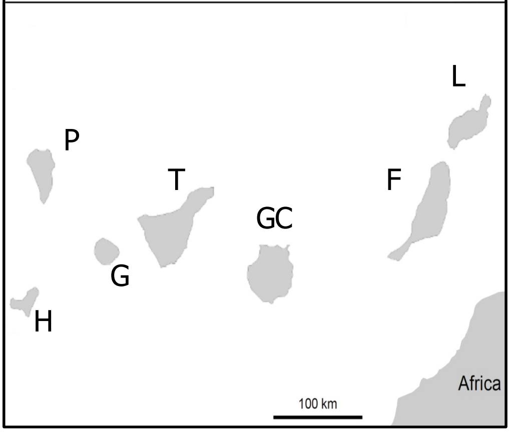
